# Short Interrupted Repeats Cassette (SIRC) ensembles of plant genomes reflects evolutionary route

**DOI:** 10.64898/2026.03.27.714674

**Authors:** Igor Vladimirovich Gorbenko, Dmitriy Yurievich Scherbakov, Karolina Mikhailovna Zverintseva, Yuri Mikhailovich Konstantinov

**Author notes:** Correspondence (I.V.G.).

## Abstract

Short Interrupted Repeats Cassettes (SIRC) are recently discovered eukaryotic DNA elements possessing many traits of satellite DNA and mobile genetic elements, and consisted of short direct repeats interspersed with diverse spacer sequences. The SIRC ensemble of individual species is highly heterogenous and cannot be studied using alignment methods. It was found that number of similar SIRC sequences in a given pair of species is in general correlated with their taxonomic distance, and, at the same time, closely related species can possess very diverged SIRC ensembles, which makes SIRC evolutionary pattern closer to mobile genetic element type. The SIRC sequences make up clusters with comparable sequence patterns, that are likely to demonstrate doublet evolutionary model which strongly supports that the SIRC structure is supported by the evolutionary selection. Several SIRC sequences of *Arabidopsis* were found to be of ancient origin with traceable evolution history as far as to the moss clade. We carried out unbiased detection of SIRC ensembles in 10 plant genomes and found that, despite very high intraspecies heterogeneity, SIRC sets possess strong interspecies phylogenetic signal.

**Key message:** Short Interrupted Repeats Cassettes are elements of ancient origin, and could potentially be used to trace organism history, and to facilitate syntheny and Hi-C analysis.

## 1. Introduction

Earlier (Gorbenko et al., 2023) we reported on the identification of novel type interspersed repetitive DNA elements – Short Interrupted Repeats Cassettes – within the nuclear genome of *Arabidopsis thaliana*. The elements are composed of interspersed DNA direct repeats interrupted by diverse spacer sequences and are structurally similar to prokaryotic CRISPR and STAR-like elements. The function of SIRC elements is yet to be determined, although their complex and non-random structure suggests some functionality. The functions of CRISPR are diverse and in general related to immunity and gene expression control (reviewed in (Koonin et al., 2017)), while STAR-like elements were recently proposed to be a regulator of translation that affects antibiotic resistance (Mediati et al., 2024). SIRC are distributed throughout *Arabidopsis* genome with maximums in pericentromeric regions, like typical mobile genetic elements. Our working hypothesis is the emergence and propagation of SIRC as complex internal structure of MITE transposons (Gorbenko et al., 2023). Similar structure had already been reported as early as 1992 to take place in *Zea mays* S3 plasmid, where it constitutes terminal inverted repeats and have two additional direct repeats in a plasmid body (Leon et al., 1992), although that time it was the only example of the structure, so it wasn’t attributed to a distinct repeat type. The role of the element in plasmid is unknown, but it could be related to ORF1 promoter sequence and the binding site of terminal proteins.

The aim of a current work is to classify SIRC sequences in *A. thaliana* genome and to conduct interspecies comparisons of SIRC ensembles in plant genomes. Using the minimum spanning trees along with alignment-free method we demonstrated that SIRC propagation is clearly connected to mobile genetic elements. We separated SIRC ensemble of *A. thaliana* into clusters and showed that these clusters have conservative consensus secondary structures using Bayesian likelihood ratio estimation. We detected SIRC unbiased set using SIRCfinder on 10 plant genomes and found that interspecies comparisons divide SIRC into two distinct types – highly conserved and diverged, which is likely to be connected to mobile genetic elements activity during new species emergence.

## 2. Results

### 2.1. Minimum-spanning trees

The Minimum Spanning Trees (MSTs) obtained are shown in Fig.1. It is interesting that in case of full SIRC sequences (Fig.1B) the tree has star-like topology, and there clear is a big central node that is formed by SIRC CP096026_89_10 possessing 118 connections with another nodes. This 49-nt length SIRC possess 2 spacers and a 10nt DR (AAGTTCTTAA) which is widespread among MGE sequence (180 occurrences detected) and is related to edge-deletion (ED) cluster 291 (of size 1) and label propagation (LP) cluster 392 (of size 136). This SIRC is overlapped with MGE AT3TE38565 of ATCOPIA65 family (in position −108 from GAG polyprotein gene start) and with sRNA smallRNA-15761 (percentage of overlapping (POL) = 100 % in both cases). According to (Oberlin et al., 2017) the AT3TE38565 is not expressed. We checked the activity of other SIRC-possessing MGEs and found that one branch of MST (full sequences) is clearly associated with MGEs that are able to be activated upon *ddm1* and/or *met1* knockout background (Supplementary Figure S1), which speaks in favor of their propagation along with MGEs. We suggest that the tree core being made of non-MGE SIRCs in general, and outer layers of the tree made of active MGEs associated SIRCs, speaks in favor of the hypothesis of the tree topology having the oldest SIRC in a center and moving forward in time to tree edges composed of young SIRC sequences.

**Fig. 1.**
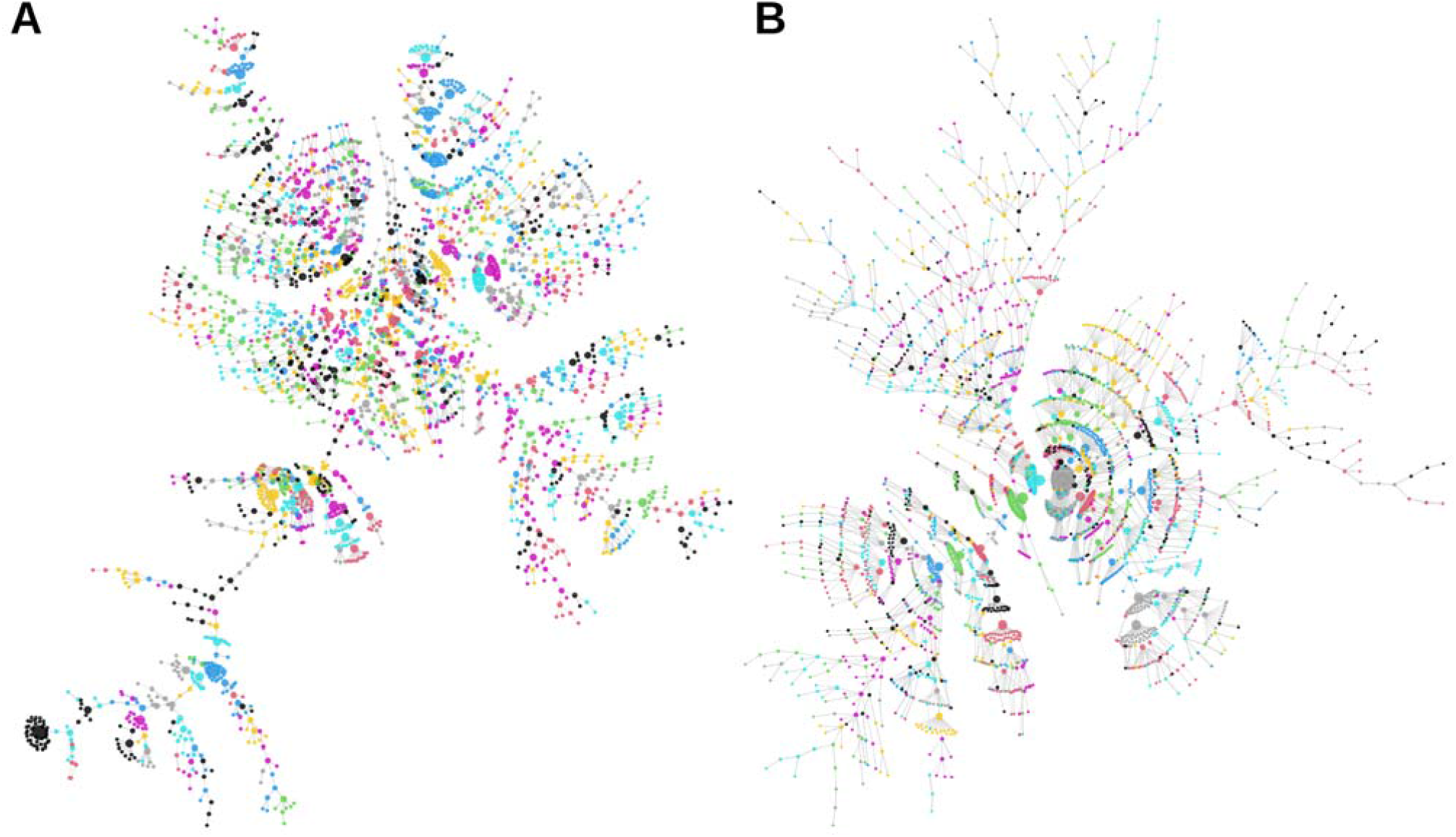
The obtained minimum-spanning trees using the DR consensus sequences (A) or full SIRC sequences (B). The nodes are colored according to label propagation community detection method.

We selected each node of both trees and calculated mean differences between the node and its closest neighbors (Supplementary Figure S2). It is clear that the most variable mean-difference parameter was len.diff – the mean-difference of lengths of SIRC of adjacent nodes, while DR length median variance was only about 2 nt, and for spacers it was of 0 nt, which speaks in favor of DR-spacer structure significance. Surprisingly the spread of all assessed parameters was a bit smaller in MST obtained from full SIRC sequences, suggesting that this approach is more promising.

### 2.2. SIRC clusterization

Using Uniform Manifold Approximation and Projection (UMAP) (McInnes et al., 2020) for dimensionality reduction of hexamer frequencies data, we estimated clusters with HDBSCAN algorithm (McInnes et al., 2017). The obtained data possessed 144 clusters of 5-81 sequences and 733 sequences were filtered out as noise points (Fig. 2A)

**Fig. 2.**
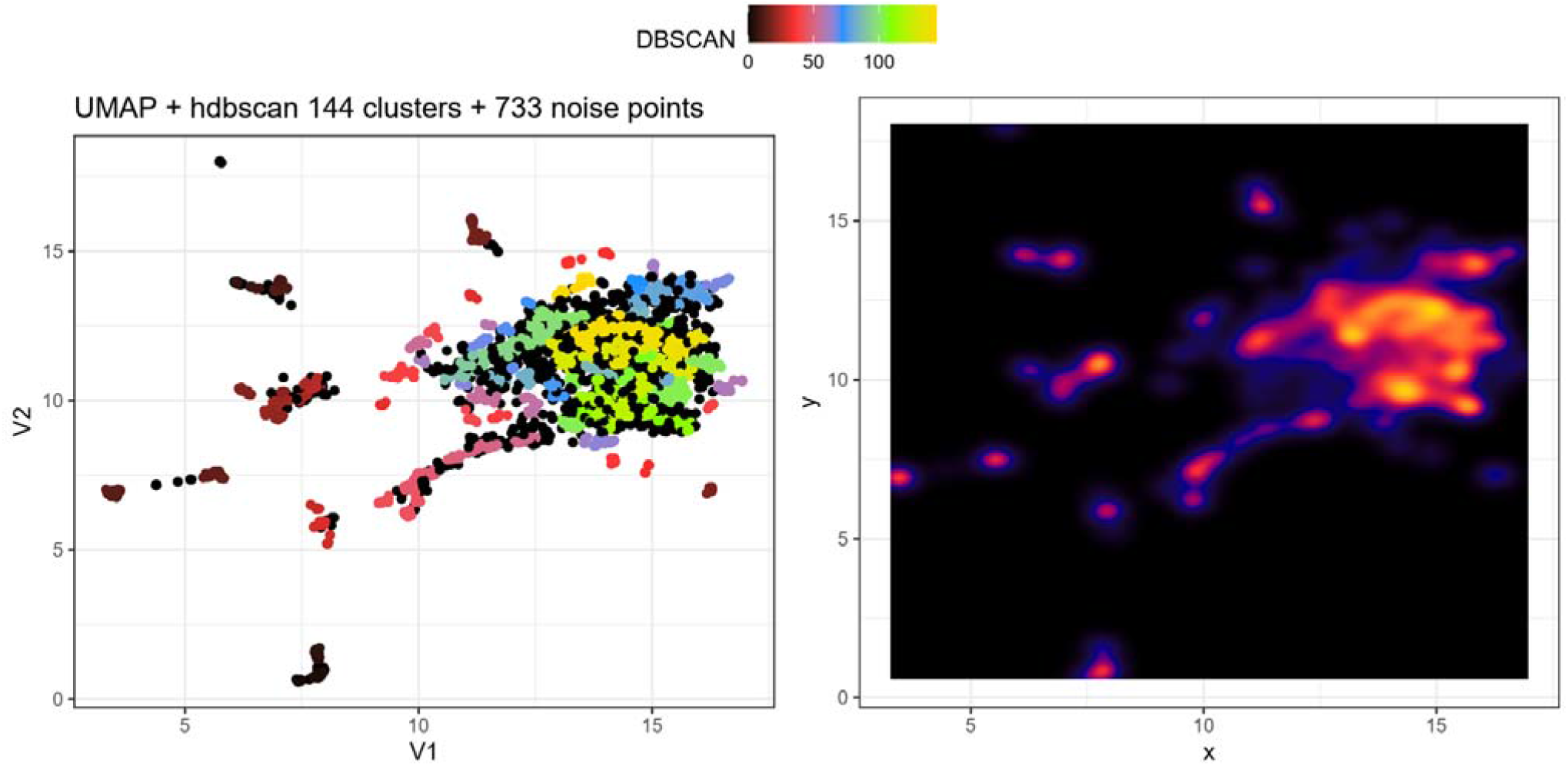
A: 2D-representation of hexamer frequencies data of SIRC with clusters estimated with HDBSCAN algorithm, noise points are colored black; B: Density map of 2D-representation of hexamer frequencies data that clarifies the heterogeneity of data point clouds.

The sequences were aligned with MUSCLE algorithm (Edgar, 2004) with subsequent run of Vienna RNAalifold (20 □) (Lorenz et al., 2011). It was found that 39 clusters (494 sequences in total) possessed consensus secondary structures (Fig. 3). Next, we utilized custom script formatting input file for MrBayes (Ronquist et al., 2012) taking into account the consensus hairpin structure revealed by Vienna RNA package as described above by assigning doublet model to the paired bases (https://github.com/dysh/STEMS) to determine if the consensus secondary structure is supported by the evolutionary selection. The method is based on the Bayesian estimation and odd ratio test to determine whether doublet evolutionary model probability is significantly different from GTR evolutionary model for the given sequence alignment. The doublet evolutionary model leads to secondary structure conservation. We found that doublet evolutionary model was significantly more probable than GTR model for 35 SIRC clusters, and for 4 clusters the likelihood ratio was higher than 30 (Fig. 4). This result supports the hypothesis of consensus secondary structure of SIRC sequences.

**Fig. 3.**
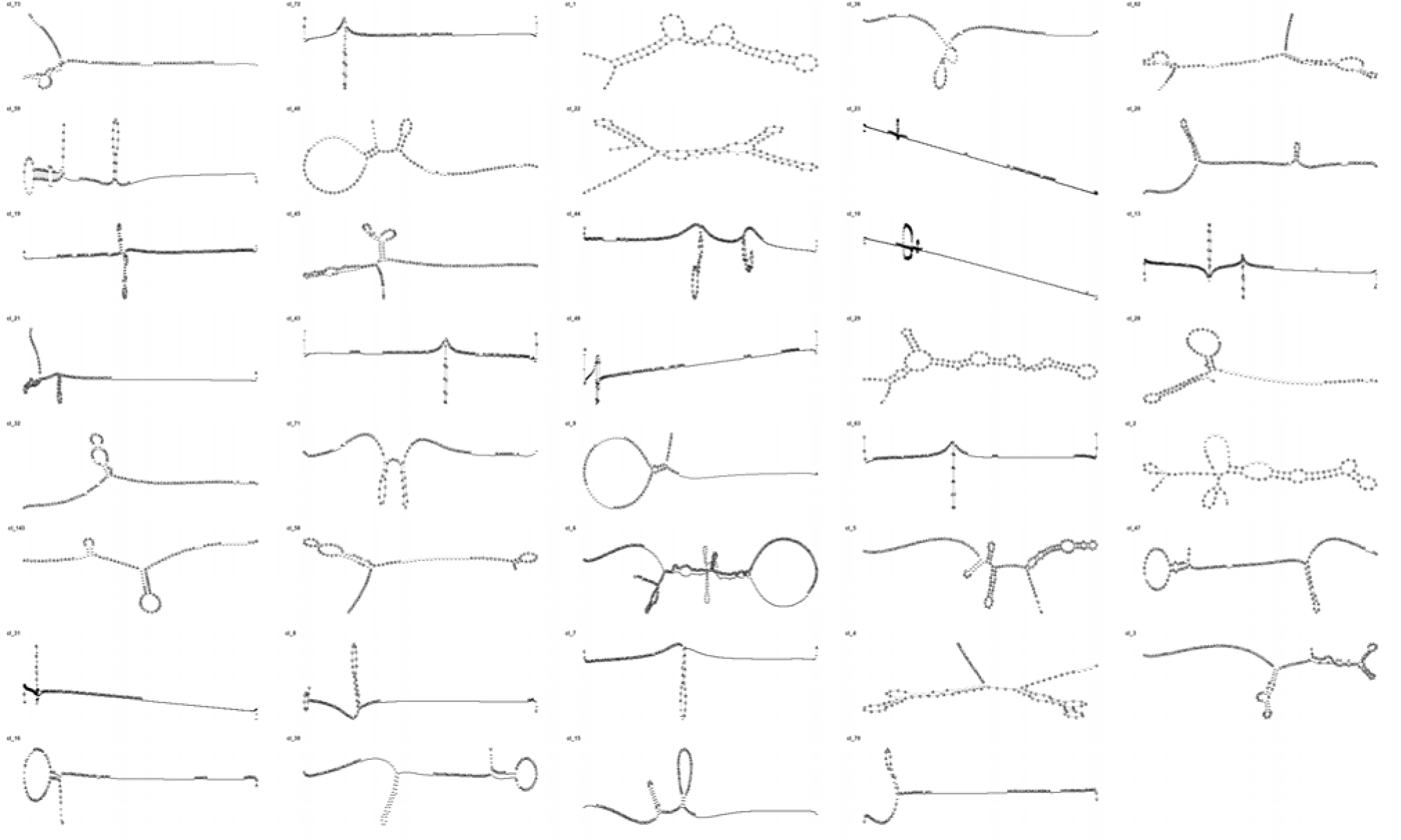
Consensus secondary structures of 39 SIRC clusters.

**Fig. 4.**
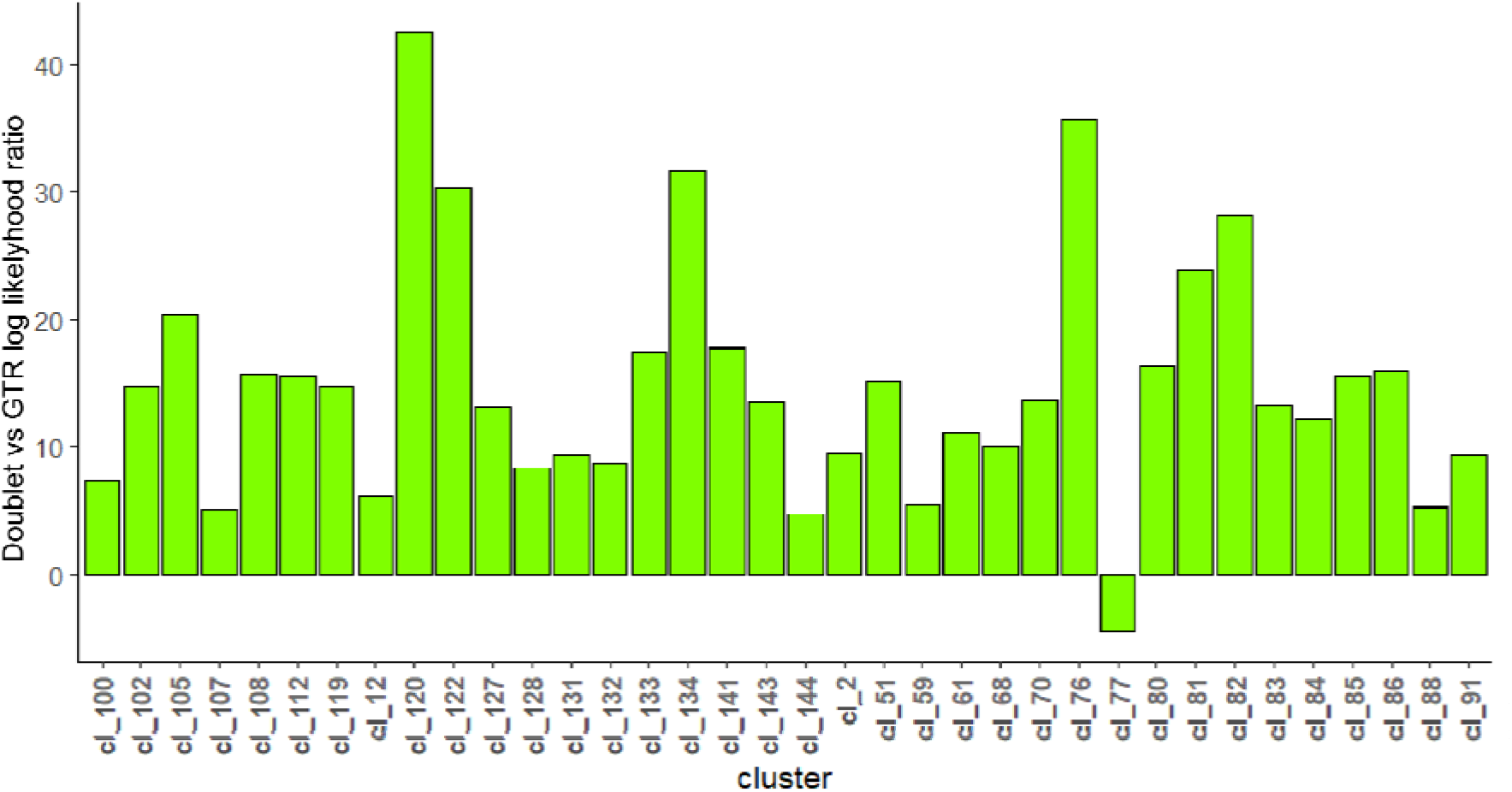
The likelihood ratio estimated for SIRC clusters.

### 2.3. SIRC family

Using BLAST, we searched for selected SIRC sequences in other higher plant species, and filtered them species-wise to leave only one similar sequence with biggest length per one selected *A*.*thaliana* SIRC. It was found that monocots do not possess similar sequences at all. Among dicots, 425 SIRC sequences were found in genome of *A. suecica*, that originated about 16000 years from the hybridization event of *A. thaliana* and *A. arenosa*, and 89 of the selected SIRC in *A. arenosa* genome (Fig. 5). Surprisingly, the number of sequences similar to selected SIRC is not strictly dependent on plant taxonomy – it is clear that the further away from Arabidopsis clade the taxon is – the fewer similar SIRC could be found, but closely related species (e.g., C. *sativa* and C. *hispida)* may possess very different numbers of similar sequences. Despite many found similar sequences, in total plants share not so many of them (Supplementary Figure S4).

**Fig. 5.**
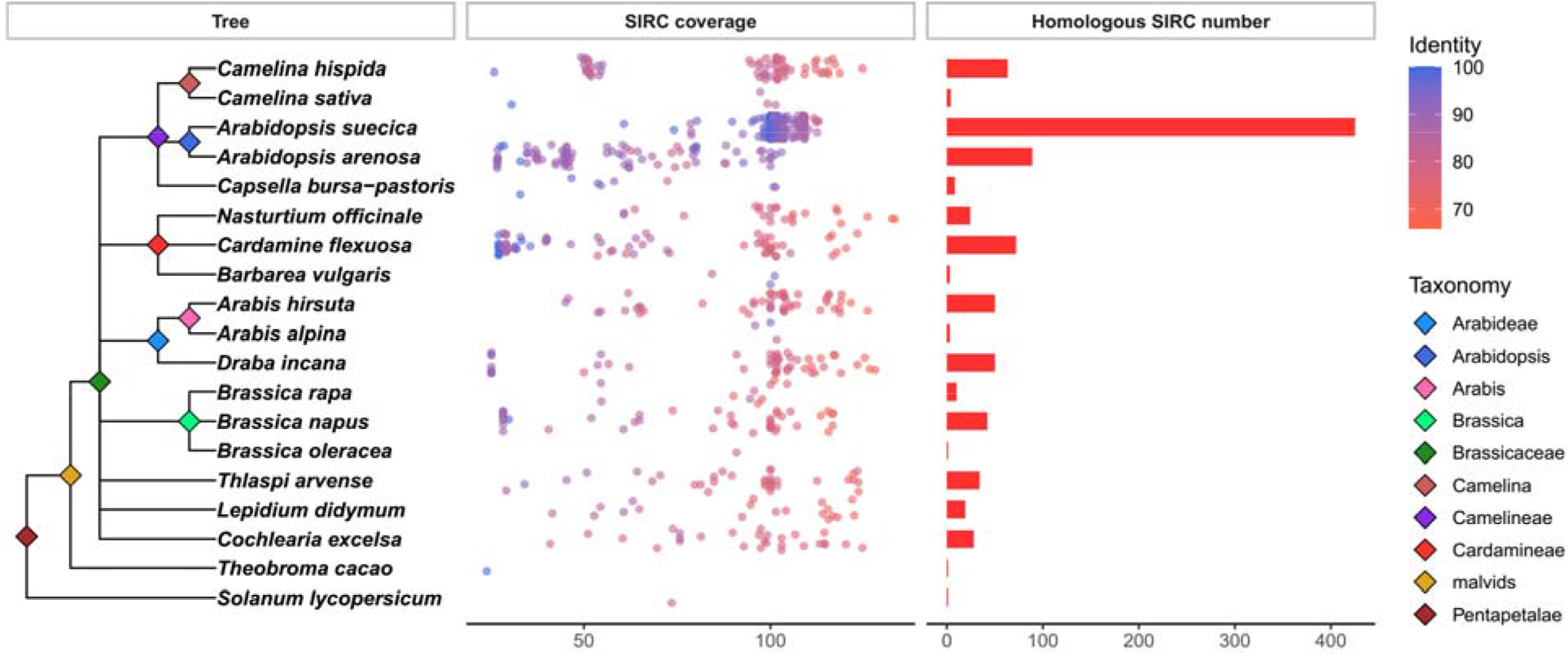
Selected SIRCs in another species genomes. The tree is based on modern plant taxonomy (https://www.ncbi.nlm.nih.gov/Taxonomy/). Point color in “SIRC coverage” section represent percentage of identity; coverage is a A. thaliana SIRC length divided by alignment length.

Using the obtained data, we selected 3 SIRCs and their best matches from ten related species, aligned with MUSCLE (Edgar, 2004) (Supplementary Figure S3), and conducted ML phylogenetic tree reconstruction with prior substitution model selection (by BIC) (Fig. 6) with phangorn (Schliep, 2011). All three SIRCs are overlapped in *A. thaliana* with MITE MGEs and two of them possess same 14-nt DR consensus (TCTATAAAITAATA). The mentioned sequences must be of ancient origin, since similar sequences were found in T. *arvense* and C. *excelsa*, and in several cases the similarity degree to *Arabidopsis* SIRC is higher than that for closely-related species. We suggest this phenomenon can be explained by alteration of MGE content that could take place in case of e.g., altered environmental conditions, leading to change in that particular MGEs (Baduel & Quadrana, 2021; Ito, 2022; Pimpinelli & Piacentini, 2020) – that’s why species closely related to *Arabidopsis* could lose SIRC similarity. Substitutions among three concatenated sequences in ten species are best explained with HKY+G(4) model.

**Fig. 6.**
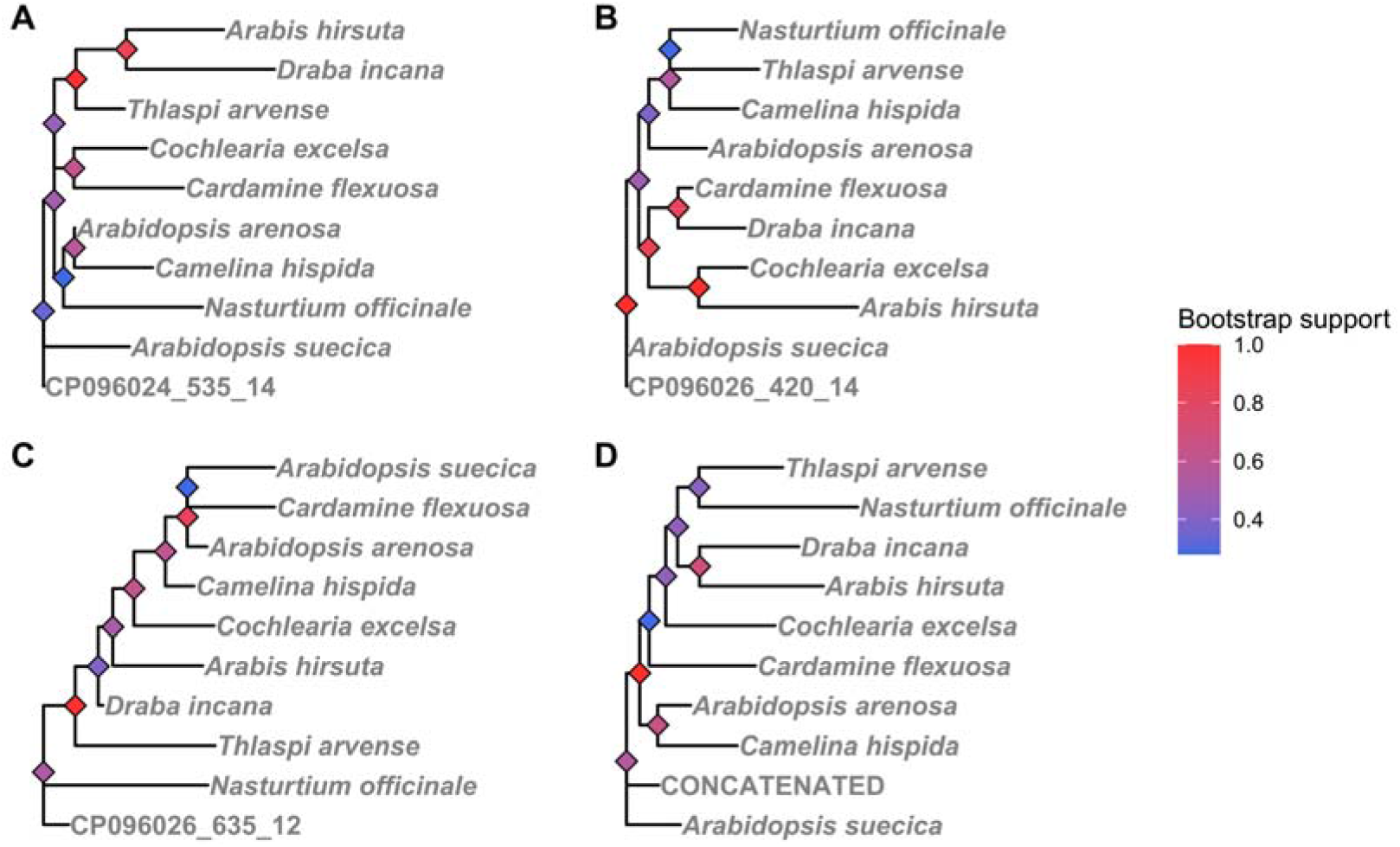
The ML phylogenetic trees obtained using SIRC-like sequences shared among ‘10 species: A - CP096024 535 14; B – CP096026_420_14; C – CP096026_635_12; D – Concatenated sequences.

We searched further for traces of the longest of selected SIRC sequences (CP096024_535_14) in plants and found 204 specie outside the *Arabidopsis* clade, and the most far were two *Sphagnum* species (S. *contortum* and *S*.*tenellum)* and *Marchantia polymorpha* subsp. ruderalis (Supplementary Figure S5). The estimated maximum likelihood tree is presented in Fig. 7.

**Fig. 7.**
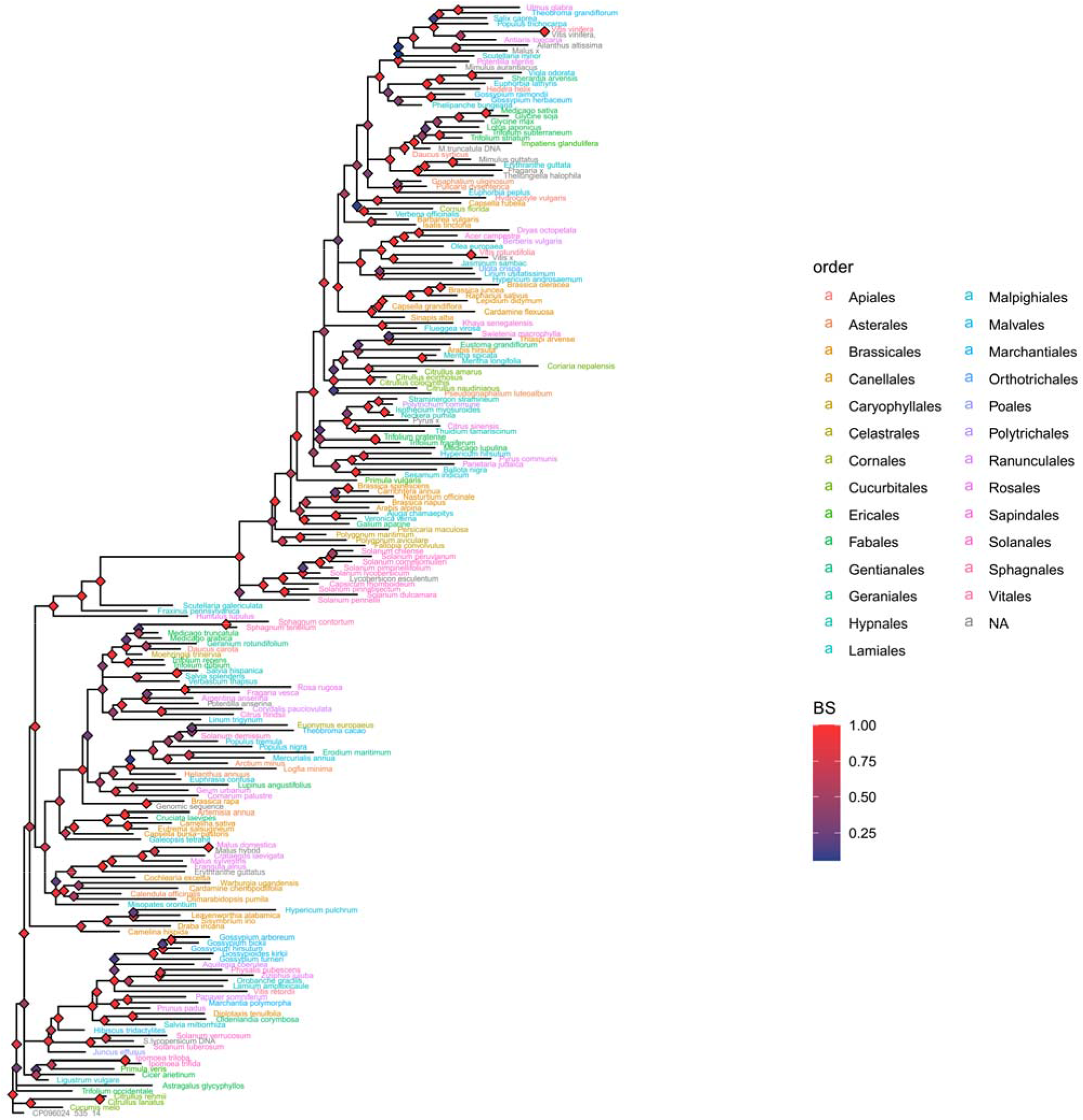
Optimal maximum-likelihood tree under the TVM+G(4) model of best matches of SIRC CP096024_535_14 among higher plants. Node color represents bootstrap support (BS).

### 2.4 Unbiased set analysis

Using the SIRCfinder R package (https://github.com/Daynoru/SIRCFinder) we scanned complete genomes of 10 species (*A. alpina, A. arenosa, A. suecica, A. thaliana, B. oleracea, Barbarea vulgaris, C. sativa, C. bursa pastoris, S. lycopersicum* and T. *cacao)* for SIRC. The obtained data was reduced via elimination of SIRC with short DR (10 nt), and analyzed using PERCON algorithm (Kazakov et al., 2003) by making comparisons of sequences octamer dictionaries with 100 random sequence pairs of same size. The minimum spanning tree obtained from fully connected graph of analyzed sequences (14338 vertices, edge weights represent PERCON r_°_ values) is presented in Fig. 8A. The numbers of intraspecies and interspecies edges in MST were 5698 and 8639, respectively. The central part of a graph is composed of similar SIRC from different species, assuming that there is a population of near-conserved SIRC, even among phylogenetically distant species. We also conducted AMOVA-like analysis on the obtained data and found 1.097 between/within distance ratio, with R^2^ = 0.856 and *P =* 0.0099 (F_stat_ = 9502.334), which allows us to assume that SIRC in general have strong phylogenetic signal.

**Fig. 8.**
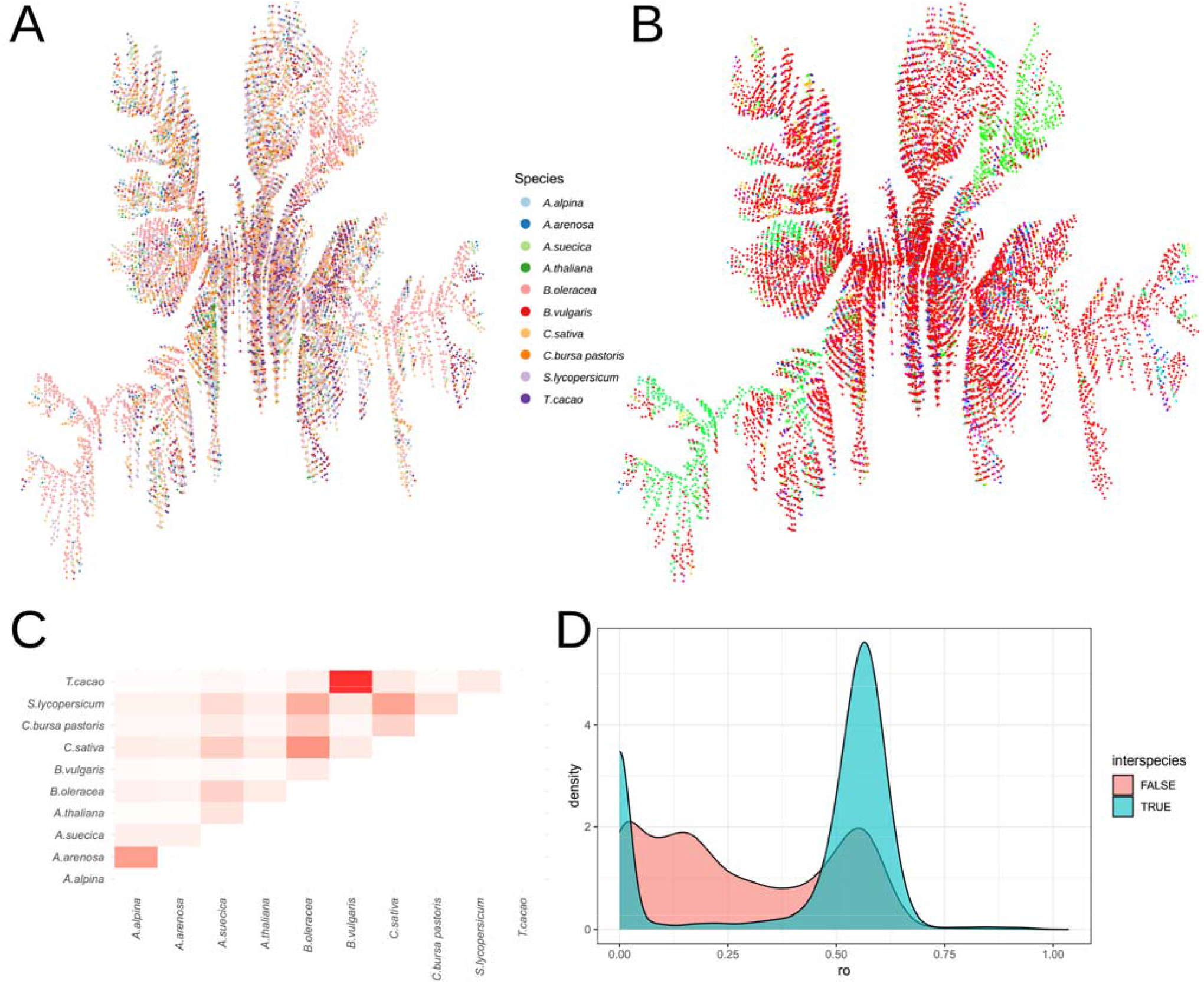
**A:** Minimum spanning tree of 14338 SIRC sequences detected in 10 species. Edge weights were PERCON r_°_ values. **B:** Communities (1534) detected in fully-connected graph via labelpropagation algorithm (Raghavan et al., 2007). **C:** A heatmap representation of interspecies MST edge numbers. **D:** Density of PERCON r_°_ values distribution among within-species and interspecies MST edges.

Using weighted label propagation algorithm (Raghavan et al., 2007) we detected 1534 communities (Fig. 8B), mostly very small in size (2-3 sequences), but one consisted of 10011 sequences (the PERCON r_°_ threshold was of 0.6), which was marked by red vertices in Fig.8B.

Many of MST edges were interspecies, indicating cases when SIRC of another species is closer in sequence than other within-species SIRCs. The adjacency matrix of SIRC interspecies connections is presented in Fig. 8C. The analyzed species are considered closely-related, except for *Barbarea vulgaris* and T. *cacao*, which had the greatest number of detected interspecies MST edges. The distribution density of inter- and intraspecies r_°_ values (Fig. 8D) shows that one of the distribution peaks of interspecies r_°_ values is around 0, the other is around 0.55, which speaks in favor of SIRC division into two groups – conserved and diverged.

## 3. Discussion

Short Interrupted Repeats Cassettes are recently described genetic elements, found and described in the genome of a model plant *A. thaliana*. Though the function of SIRC remains unknown, complex internal structure and results of the current work allow us to suggest functional significance of these elements. In the current work we conducted complex analysis of SIRC elements in 10 plant species using combination of bioinformatics and evolutional biology methods.

Using minimum spanning trees approach we found that within-species SIRC propagation is likely to be due to mobile genetic elements activity (Fig. 1, Supplementary Figure S1), which is in a good agreement with our previous results (Gorbenko et al., 2023). The co-option of a transposable element as a shuttle for other genetic elements propagation is known for different domains of life. For example, a part of C. *elegans* germline-specific promoters were co-opted from CERP2 and CELE2 transposons of MITE family (Carelli et al., 2022). In human, some transposons are utilized as enhancers that are active in cancers of endodermal lineage (Karttunen et al., 2023). Also, it is thought that, in human, transposons located in immune-related loci facilitate active selection and functionalization of transposon motifs, which may have led to accelerated evolution of enhancers in immune cells (Ye et al., 2020). In mouse, large number of LTR transposons facilitates proper execution of tissue-specific gene expression programmes (Todd et al., 2019). Transposable elements can generate transcription factor binding sites (TFBS) that in some cases affect chromatin 3D organization and affect regulation of distal genes expression (Avramova et al., 1998; Choudhary et al., 2020; Diehl et al., 2020).

Using UMAP with HDBSCAN cluster detection we were able to filter out highly-diverged “noise” SIRC and isolate SIRC clusters (Fig. 2). Several clusters had consensus structural patterns that we studied via estimation of consensus RNA secondary structure (Fig. 3), and MrBayes software allowed us to estimate log likelihood ratio for evolutionary models of these structures (Fig. 4). The results obtained shows that the doublet code was more likely for most of the found clusters, indicating that the compensatory model describes these clusters well, thus the base pairings conservation is likely to be supported by the selection (Fig. 4). The compensatory evolution model is known for such structures as rRNA, tRNA, ribonuclease P RNA and other, where the base-pairings form structures crucial for RNA functions (Higgs, 1998; Tillier & Collins, 1998), and for RNA-RNA interacting pairs (Durand & Storz, 2010; Kern & Kondrashov, 2004; Knies et al., 2008; Schaufele et al., 1986; Waters & Storz, 2009).

Using BLAST software, we found sequences similar to *A. thaliana* SIRC in another species and observed a distinct trend – the number of similar sequences in general decreases with taxonomic distance increase from *Arabidopsis* to another species, while closely related species may possess very different numbers of similar sequences, indicating that SIRC evolution is close to mobile genetic elements type (Fig. 5). This is in a good agreement with up-to-date data on novel species formation – extreme environmental changes promote severe genetic stress, which increases probability of organism adaptation due to higher mutation rate (Baduel & Quadrana, 2021; Carfora et al., 2025; Heinen et al., 2025; Ito, 2022; Pang et al., 2021; Pimpinelli & Piacentini, 2020; Ramos-Onsins & Ferretti, 2026). Similarly, different stress amount leads to different duration and strength of mobile elements activity (Miryeganeh & Armitage, 2025; Pimpinelli & Piacentini, 2020), which can be traced from SIRC differences (Supplementary Figure S1). On the other hand, similar to SIRC sequences can be very conserved in distantly related species, indicating the ancient nature of these elements (Fig. 6, Fig. 7).

Using the SIRCfinder R package we detected SIRC in 10 plant genomes. It was found that SIRC in general possess very strong Phylogenetic signal (R^2^ = 0.856, *P =* 0.0099, F_stat_ = 9502.334), despite relatively low between/within species distance ratio of 1.097. This fact emphasizes that despite persistent overdispersed “noise” in different species SIRC ensembles, the evolutionary history of SIRC is firmly encoded in their presence/absence in particular loci of different species. The distribution of SIRC distances indicate that there are mainly two SIRC classes during interspecies comparisons – highly conserved (r_°_ ~ 0) and dispersed (r_°_ ~ 0.55), which additionally makes SIRC closer to mobile genetic elements type of evolution (Fig. 8D). The key for solvation of this evolutionary paradoxus is likely to be the importance of SIRC structure consisted of direct repeats interrupted with diverse spacers sequences, which is the one under the natural selection and could potentially be encoded by thousands of different sequences.

The obtained data shows that SIRC are dynamic genetic elements, and their evolution and conservation is tightly bound to mobile genetic elements. It’s likely that stable SIRC are tightly bound to their genetic positions even comparing different species, which emphasizes the perspective of SIRC elements usage for Hi-C sequencing of syntheny analysis.

In future works the usage of SIRCfinder on hundreds of species will allow to trace the evolution of SIRC structural motif and find out if it correlates with the key events in plant genome evolution like water-to-land plants conversion or flowering emergence.

## 4. Materials and Methods

The principal workflow of the first part current paper is presented in Fig.9.

**Fig. 9.**
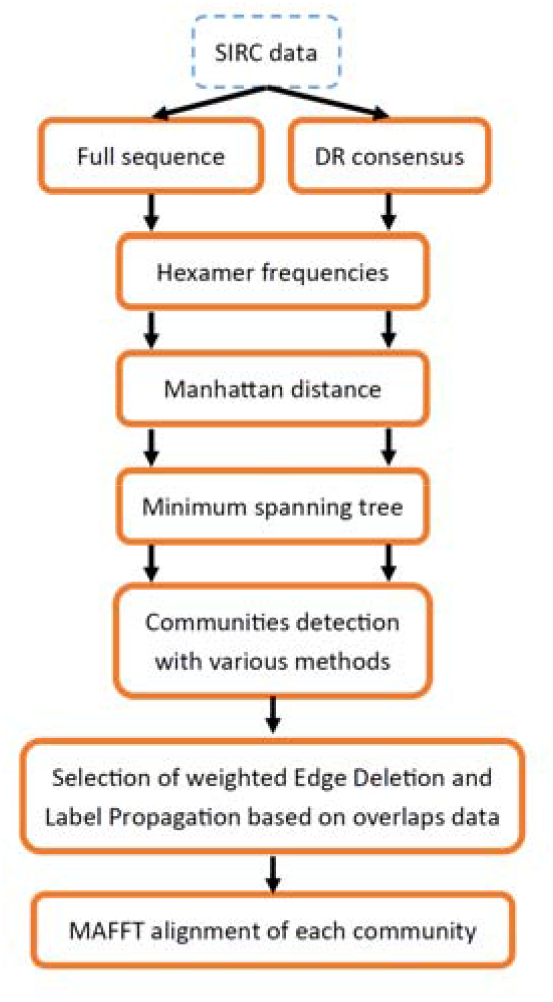
The principal workflow of the first part of the current work.

### 4.1. SIRC clusterization

The full SIRC sequences were extracted from *A. thaliana* genome. The following workflow was applied to both full sequences and DR consensuses. As SIRC are extremely diverse and unalignable in total, for initial clustering we used hexamer frequencies extracted via Biostrings R package (Pagès et al., 2025), and calculated pairwise Manhattan distances using parallelDist R package (Eckert, 2022). Next we used genieClust package (Gagolewski, 2021) to estimate minimum spanning tree (MST) using weighted Jarnik (Prim/Dijkstra)-like algorithm (Cheriton & Tarjan, 1976). To divide the MST into clusters, we deleted all edges with weights more than a threshold weight, estimated using validation via the Ball-Hall index (Ball & Hall, 1965).

### 4.2. SIRCfinder R package

The summary of main SIRCfinder algorithm is presented in Fig. 10. The package function SIRCFinder() is the full C++ implementation of algorithm, powered by Rcpp. The algorithm includes 7 main parts. First, the input sequences are separated by chunks (1Mbp by default). Then the 1^st^ part “Cores detection” performs parallel search of imperfect (80% identity by default) direct repeats in a sliding window (2000 kb by default). The algorithm bypasses soft masked regions as well as low complexity regions. The found core groups, or protocassettes, are separated to insure proper spacer-DR size proportions. In case of overlaps – the protocassette with best entropy weight is chosen. In cases of perfect core groups overlapped (W = 1) – heuristic algorithm performs merge of protocassettes DRs resulting in extension of DR lengths. The 2^nd^ part “Cores extension” detects proper DR-spacer boundaries by scanning the protocassette for rapid Shannon entropy jump that occurs in spacer start position. The 3^rd^ part “Splitting” performs protocassette splits in cases of improper-sized spacers. Sequentially all resulting protocassettes of improper DR number are being deleted. The 5^th^ part “Refinement” performs additional genome analysis to add missed DRs to protocassettes. The 6^th^ and 7^th^ parts are filters that compares spacers to DR consensus and spacers with spacers respectively to ensure no tandem repeats passed through the pipeline by mistake. The result is R List containing DR consensus, DR length, genomic positions of DR start, DR sequences and scanned sequence name.

**Fig. 10.**
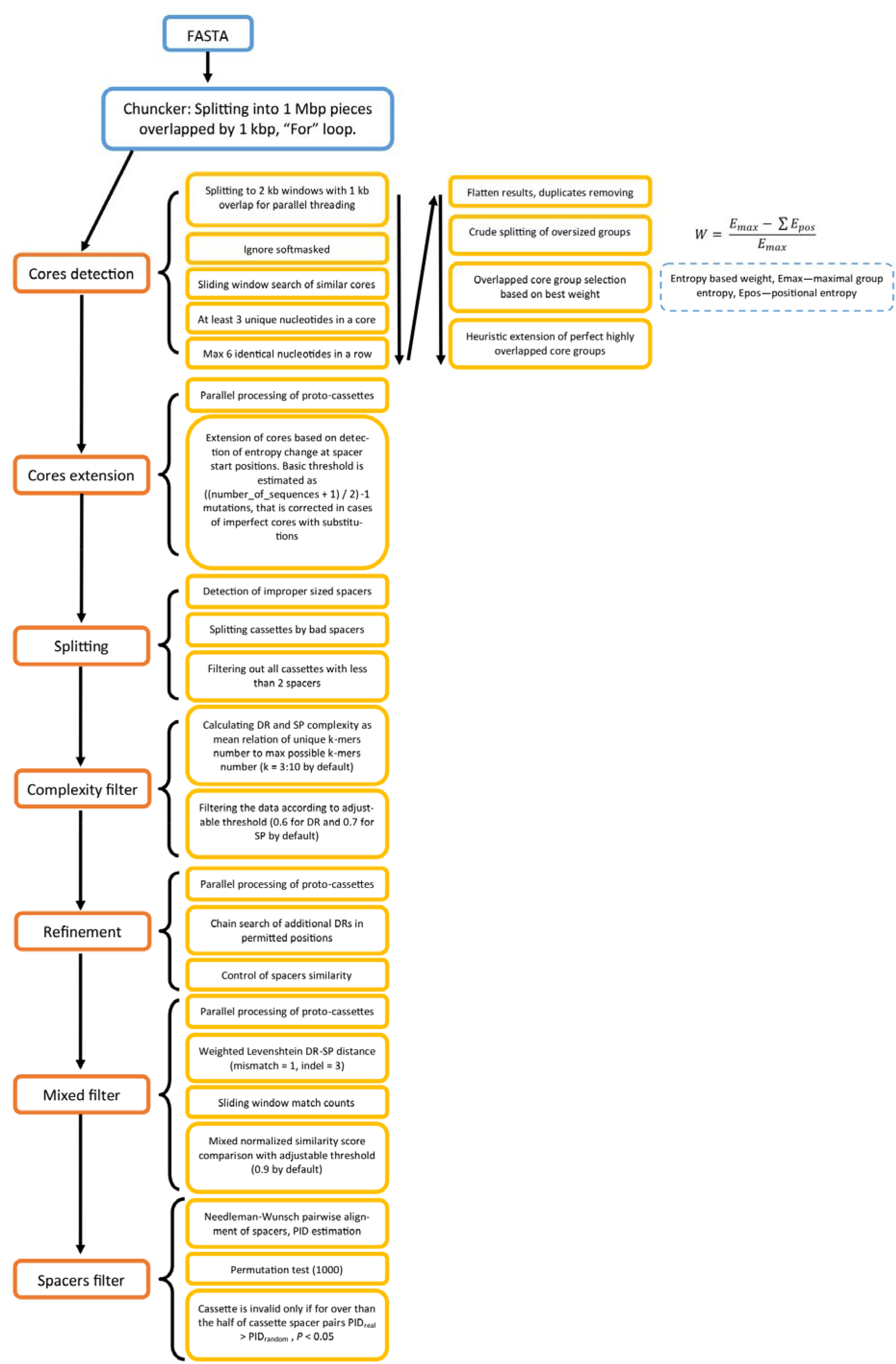
The summary of main SIRCfinder function algorithm.

The function percon_analysis() takes fasta as input and performs full analysis on SIRC pairwise comparisons by calculation of PERCON r_°_ values, formation of a fully-connected graph with label propagation algorithm community detection, minimum-spanning tree construction of a graph, and in cases of interspecies comparisons performs (argument interspecies = TRUE) performs AMOVA-like analysis calculating F-statistics, *P*-value and *R*^*2*^ to score the interspecies phylogenetic signal strength. For interspecies analysis, the sequence names in fasta file must contain species names and sequence names separated with “_”

The SIRCfinder package was tested on Ryzen 5 3600X with B550 motherboard chipset and 48 Gb RAM – the analysis for 10 genomes presented in a current study took about 3 hours of time and near 16 Gb of RAM usage. The authors want to emphasize that SIRCFinder() function of SIRCfinder R package is suited for assembled genomes, not for scaffolds or raw reads data analysis – since high number of sequences in input fasta file may cause the chunk number estimation to fail.

## Statements & Declarations

### Funding

This work was supported by the Ministry of Education and Science of the Russian Federation (project numbers 0277-2025-0001 (125021702323-2) and FWSR-2026-003 (1023032700327-0-1.6.8)).

### Competing Interests

The authors have no relevant financial or non-financial interests to disclose.

### Author Contributions

Conceptualization: I.V.G., Y.M.K.; methodology: I.V.G. and D.Y.S.; software: I.V.G., D.Y.S. and K.M.Z.; validation: I.V.G. and Y.M.K.; formal analysis: I.V.G.; investigation: I.V.G.; data curation: I.V.G.; writing—original draft preparation: I.V.G.; writing—review and editing: I.V.G., Y.M.K. and D.Y.S.; visualization: I.V.G.; supervision: Y.M.K.; project administration, Y.M.K.; All authors have read and agreed to the published version of the manuscript.

### Data availability

The R and C++ **code** used for analysis is available at **https://github.com/Daynoru/SIRCFinder;** Initial data used for analysis was the Supplementary data S5 and S6 of (Gorbenko et al., 2023).

## Acknowledgments

We thank Prof. D. B. Sloan, Prof. J. P. Mower and Dr. D. A. Afonnikov for the interest in our study and valuable advices.

## Notes

https://github.com/Daynoru/SIRCFinder

### Competing Interest Statement

The authors have declared no competing interest.

