## Supplementary Figures S1-S5 for "Short Interrupted Repeats Cassette (SIRC) ensembles of plant genomes reflects evolutionary route"

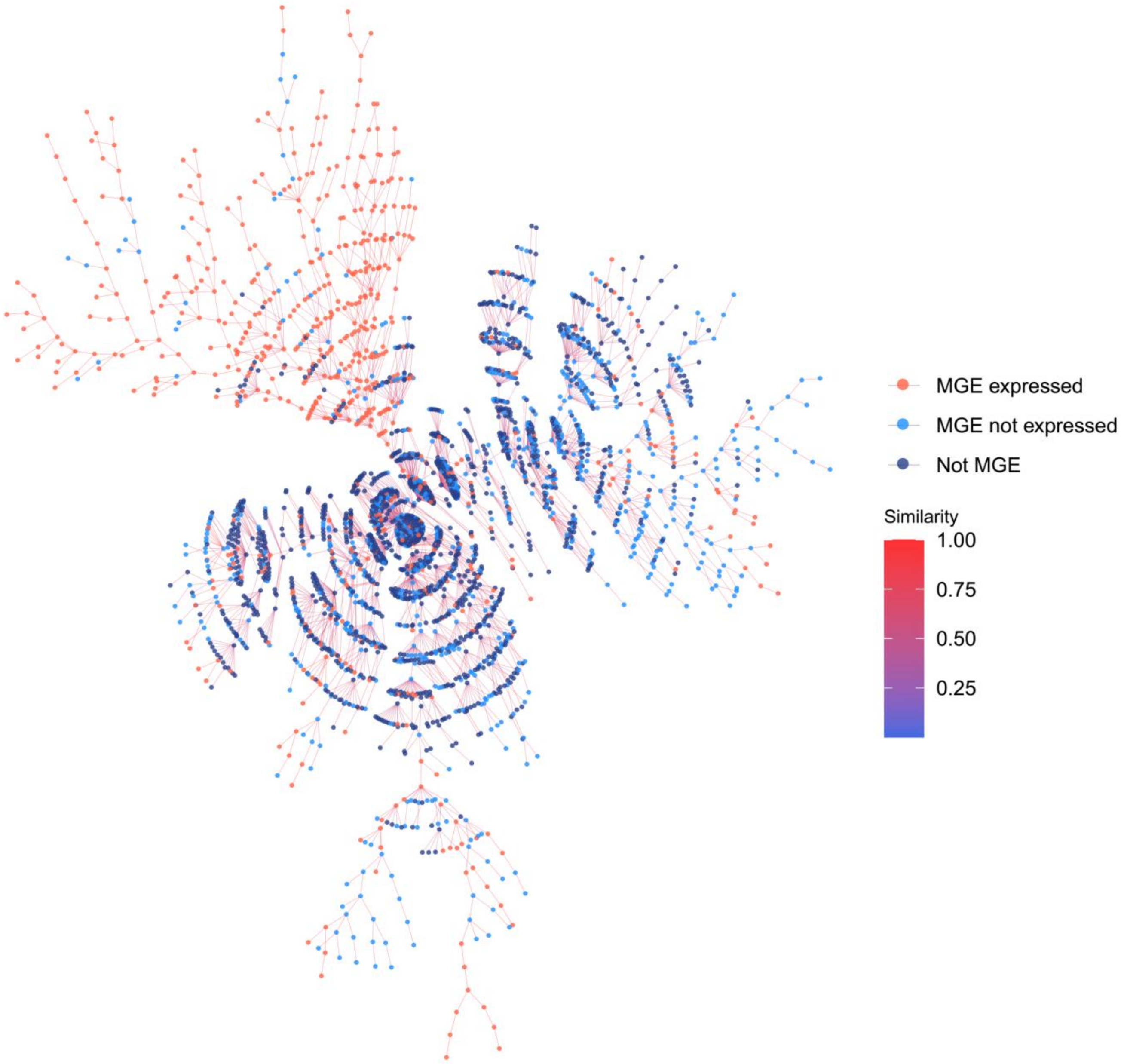


**Fig. S1.** The MST constructed using full sequences with node colors according to presence in MGE and activity of that MGE in *ddm1* and/or *met1* knockout lines (the data on MGE activity is from Oberlin et al., 2017)**.** Edge colors represent SIRC similarity (inverted Manhattan distances between hexamer frequences).

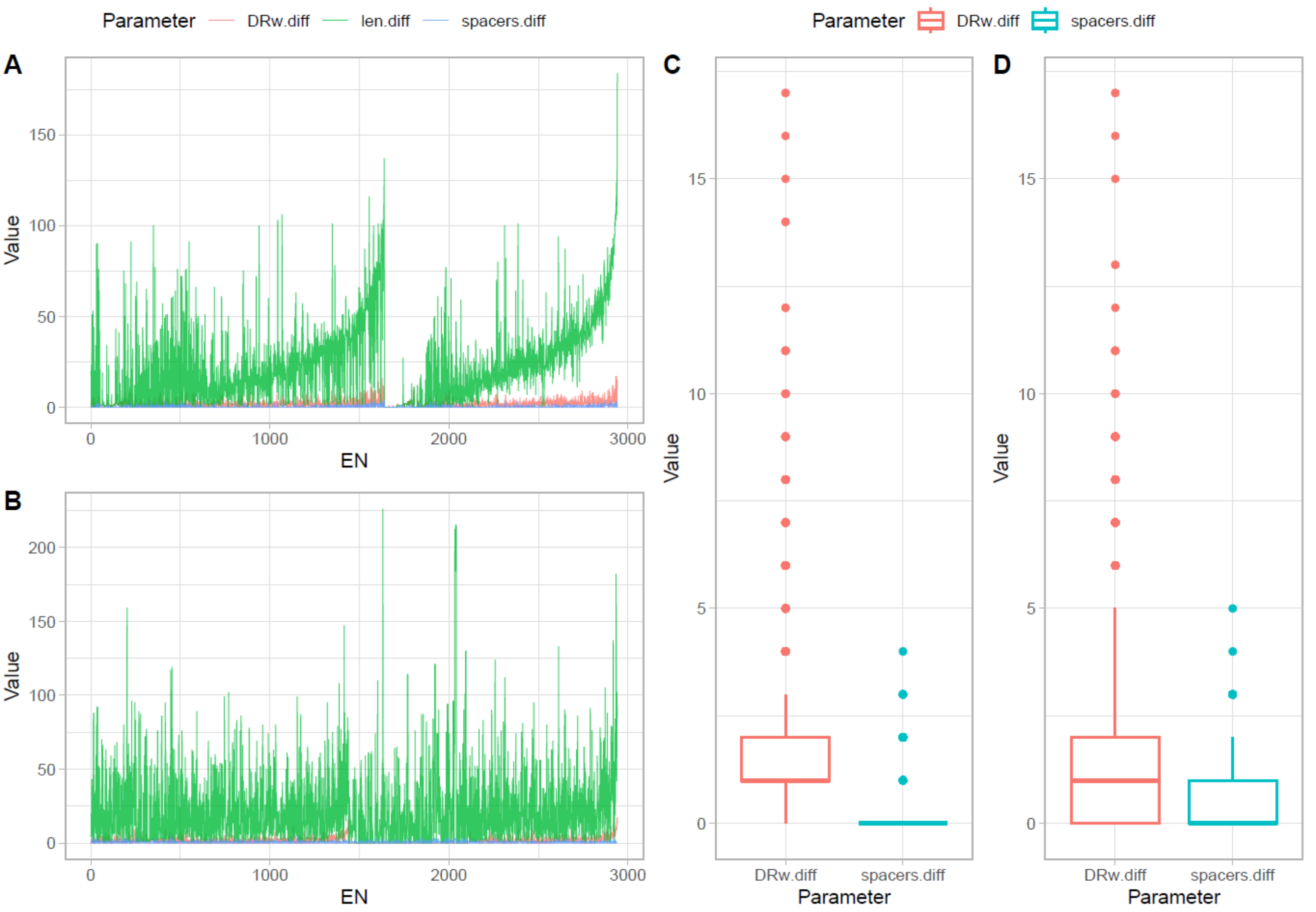


**Fig. S2.** The mean differences between each tree node and its closest neighbors. A and C – for tree obtained from full SIRC sequences, B and D – for tree obtained from DR consensus sequences only. A and B presents the mean differences across each node, C and D – the total mean difference stats. EN stands for Edge-group Number.

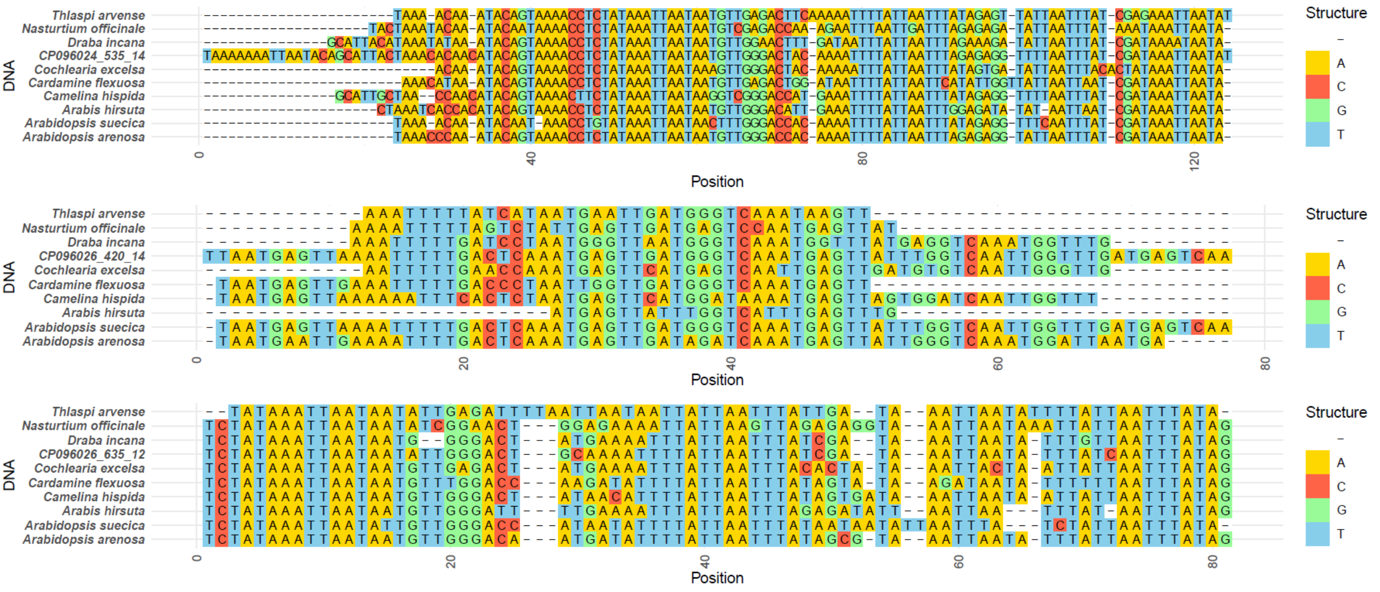


**Fig. S3.** The alignments of 3 selected SIRC sequences of *A. thaliana* and their best matches in another species.

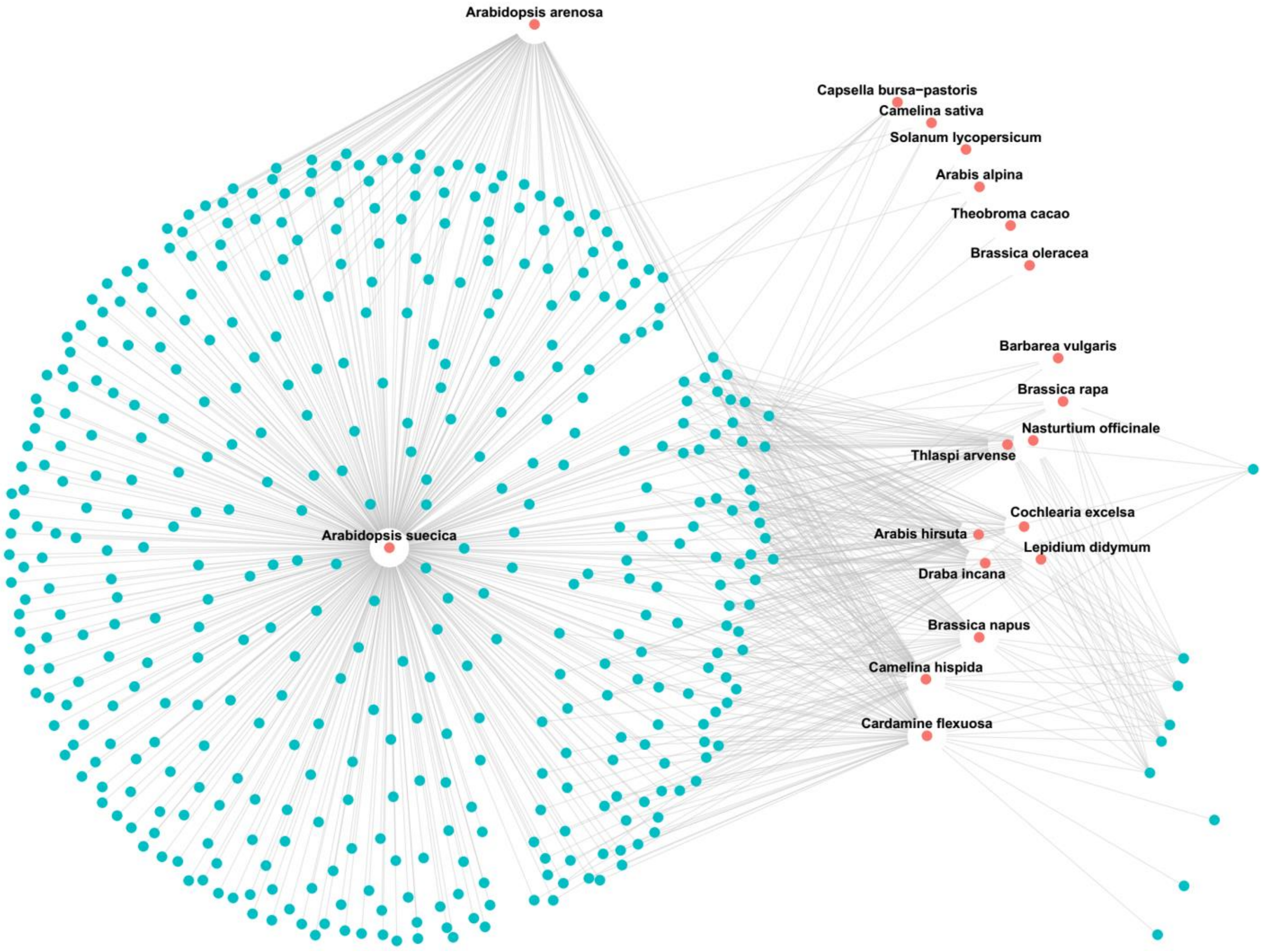


**Fig. S4.** The network of sequences similar to *A.thaliana* SIRC in selected species. *A. thaliana* sharing all the connections is not shown. SIRC are shown as blue dots, plants – as red dots.

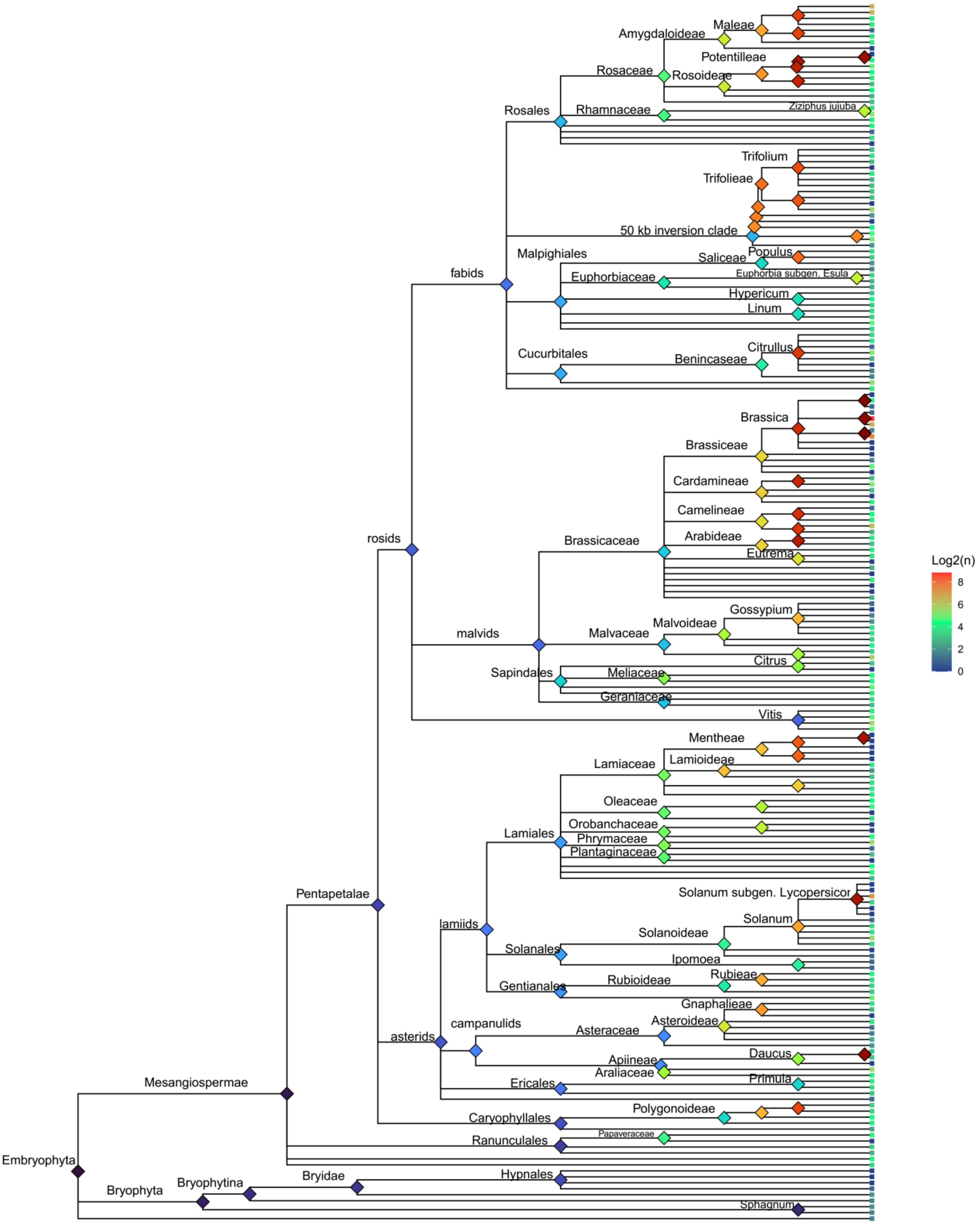


**Fig. S5.** The phylogenomic tree represented taxonomic relationships between plants among which sequences similar to SIRC CP096024_535_14 was found. The tip point colors represent logarithmic number of found BLAST hits.
